# *WUSCHEL-*dependent chromatin regulation in maize inflorescence development at single-cell resolution

**DOI:** 10.1101/2024.05.13.593957

**Authors:** Sohyun Bang, Xuan Zhang, Jason Gregory, Zongliang Chen, Mary Galli, Andrea Gallavotti, Robert J. Schmitz

## Abstract

WUSCHEL (WUS) is transcription factor vital for stem cell proliferation in plant meristems. In maize, *ZmWUS1* is expressed in the inflorescence meristem, including the central zone, the reservoir of stem cells. *ZmWUS1* overexpression in the *Barren inflorescence3* mutant leads to defects in inflorescence development. Here, single-cell ATAC-seq analysis shows that *ZmWUS1* overexpression alters chromatin accessibility throughout the central zone. The CAATAATGC motif, a known homeodomain recognition site, is predominantly observed in the regions with increased chromatin accessibility suggesting ZmWUS1 is an activator in the central zone. Regions with decreased chromatin accessibility feature various motifs and are adjacent to *AUXIN RESPONSE FACTOR* genes, revealing negative regulation of auxin signaling in the central zone. DAP-seq of ZmWUS1 identified the TGAATGAA motif, abundant in epidermal accessible chromatin compared to the central zone. These findings highlight ZmWUS1’s context-dependent mechanisms for stem cell maintenance in the inflorescence meristem.

## INTRODUCTION

In the inflorescence meristem, the delicate balance between self-renewal and differentiation of stem cells is tightly controlled by the transcription factor WUSCHEL (WUS) ^1^. This stem cell maintenance involves a negative feedback loop where WUS promotes *CLAVATA3* (*CLV3*) expression, and CLV3 represses WUS ^2^. In Arabidopsis, mutant studies revealed the functional interplay between these two components: in *clv3*, *WUS* expression expands, whereas in *wus*, *CLV3* expression markedly decreases ^3^. This feedback loop ensures a balanced stem cell population, where an overabundance of WUS increases stem cells and its scarcity depletes them ^4^. The CLV-WUS regulatory pathway is conserved among plant species, including tomato, maize and rice ^5–7^.

In maize, *ZmWUS1,* a co-ortholog of Arabidopsis *WUS,* is expressed in cells that neighbor the stem cells within the female inflorescence and the axillary meristems, which eventually give rise to mature ears ^8^. These include spikelet-pair meristems, spikelet meristems, and floral meristems, all of which contribute to kernel formation ^9^. *Barren inflorescence3 (Bif3)* mutants, overexpress *ZmWUS1* and displays small, spherical ears with a reduced kernel count, indicative of inflorescence meristem disruption ^8^. Similar to Arabidopsis, *ZmCLE7*, a functional homolog of *CLV3*, is implicated in the regulation of inflorescence growth ^10^. Diminished *ZmCLE7* expression increases kernel count, producing kernels that are narrower, less round, and deeper, whereas *ZmCLE7* upregulation lowers yield of the female inflorescence ^11^.

The influence of WUS on *CLV3* regulation is cell-context dependent. In Arabidopsis, *CLV3* is expressed in the three uppermost cell layers, L1, L2, and L3, where the stem cells are located ^12^. *WUS* is expressed in cells starting from the L3 layer and extending to more inner layers; the region where *WUS* is expressed is known as the organizing center ^13^. Although *WUS* is expressed in the organizing center that is beneath the stem cell niche, WUS protein accumulates across a wider area including *CLV3* expressing stem cells ^14^. It is posited that WUS diffuses to activate *CLV3* in these stem cells, as evidenced by the observation that a decrease in *WUS* expression leads to a reduction in *CLV3* expression ^15^. If WUS acts as an activator regardless of its cell type, it should activate *CLV3* in the organizing center cells. However, the exclusive expression of *WUS* and absence of *CLV3* expression in the organizing center suggests that WUS functions as a repressor in the organizing center ^16^. The GRAS transcription factors HAIRY MERISTEMs (HAMs) function as co-factors and their interaction suggests a model where WUS-HAM complexes in the organizing center inhibit *CLV3* transcriptional activation ^16^. In contrast, another model proposes that WUS forms homodimers that repress *CLV3* in the organizing center, but as it diffuses to nearby cells its concentration diminishes and transitions to function as a monomer that activates *CLV3* ^17^. These findings collectively highlight that WUS’s function varies dependent on cellular context, emphasizing the need for cell-specific data to fully understand its regulatory roles.

WUS is homeodomain transcription factor, that has a helix–turn–helix DNA-binding domain that targets specific *cis*-regulatory elements ^13^. There have been significant efforts to identify WUS-dependent *cis*-regulation at *CLV3* and other targets. For example, Electrophoretic Mobility Shift Assays (EMSA) and Chromatin Immunoprecipitation followed by quantitative PCR (ChIP-qPCR) have detected binding of WUS to TAAT motifs in regions flanking *CLV3*, both upstream and downstream ^14,17^. This TAAT motif is bound by other homeodomain transcription factors ^13,18^. WUS and SHOOT MERISTEMLESS (STM) also form a dimer that interacts with the TGACA motif upstream of *CLV3* ^19^. Additionally, ChIP-qPCR has revealed WUS binding upstream of the *ARABIDOPSIS RESPONSE REGULATOR 7* (*ARR7*) ^20^, whereas EMSA has identified WUS binding downstream of the *AGAMOUS* (*AG*), corresponding to a CC(A/T)6GG ^21,22^ or TTAATGG motif ^23^. Genomic approaches have also identified WUS targets. For example, ChIP-seq using inducible WUS and WUS DNA affinity purification sequencing (DAP-seq) ^24^ in Arabidopsis have identified the TGAATGAA motif ^24,25^. Microscale thermophoresis has demonstrated a preference for the TGAATGAA motif over TAAT ^26^. The crystal structure of the WUS homodimer suggests it prefers the TGAATGAA motif ^26^. These diverse and occasionally conflicting WUS binding motifs reflect the hierarchical nature of protein:DNA interactions, likely revealing that some sequences are favored over others in certain cellular contexts. This implies that its binding, as well as the specific cell types in which different co-factors or transcription factors exist, could influence WUS activity and targeting of *cis*-regulatory elements.

Although the role of *ZmWUS1* is significant in maize inflorescence development ^8^, the detailed mechanisms, and particularly its target *cis*-regulatory elements, are unknown. The complex regulatory dynamics of ZmWUS1 and its unique cell-type-specific expression pattern complicate target gene identification. To accurately identify *cis*-regulatory elements influenced by ZmWUS1 at single-cell resolution, we used single-cell Assay for Transposase-Accessible Chromatin with sequencing (scATAC-seq) using immature maize ears in wild type (WT) and the *Bif3* mutant background. By contrasting chromatin accessibility variation in WT and *Bif3* across various cell types within the developing inflorescence, we identified cell-type-specific mechanisms of *cis*-regulation and how *ZmWUS1* overexpression affects development. This research has led to a model whereby ZmWUS1 targets distinct cis-regulatory elements depending on the cell type.

## RESULTS

### scATAC-seq captures central zone nuclei in the immature maize ear

We hypothesized that overexpression of *ZmWUS1* alters *cis-*regulatory element activity in *Bif3* by binding cis-regulatory elements that are not typically targeted in WT. To assess this, we compared the chromatin accessibility landscape across cell types between WT and *Bif3* immature ears. We performed scATAC-seq in developing maize inflorescences in WT and *Bif3* using two biological replicates. After filtering nuclei based on Tn5 insertion number per nucleus, transcription start site enrichment, organellar ratio, FRiP scores, and doublet scores, an average of 4,426 nuclei per replicate remained, each with an average of ∼25,000 Tn5 insertions. Separate cell annotations for WT and *Bif3* identified 13 and 14 distinct nuclei clusters, respectively, with sub-clustering further refining cell-type annotations (Figure S2A and B; Table S1-3).

Cluster annotation was determined using gene body chromatin accessibility for marker genes (Figure 1A). Although Arabidopsis *WUS* is expressed in the organizing center, specifically in the L3 layer and more internal cells at the center of the meristem ^27^, the expression domain of ZmWUS1 in maize female inflorescence spans a broader area, covering cell layers 1 through 10 across the entire meristem, not just the middle ^8^. Furthermore, the distinction between the stem cell niche and cells containing ZmWUS1 is less defined in the inflorescence meristem, where *ZmCLE7* is expressed in the outermost cells—also the location of *ZmWUS1* expression. This pattern continues in the axillary meristem cells, where the stem cell marker gene *ZmCLE7* and *ZmWUS1* exhibit precisely overlapping expression patterns ^8^. In the *Bif3* mutant, this overlap of *ZmWUS1* and *ZmCLE7* expression extends to both the axillary and inflorescence meristems. This leads us to designate *ZmWUS1* and *ZmCLE7* expressing nuclei as the central zone that is illustrated in the accompanying sketch (Figure 1B). *ZmWUS1* gene body chromatin accessibility was widespread and enriched in WT Cluster (C) 3-1 & 3-3, and in *Bif3* C14 (Figure 1C). *ZmCLE7* exhibited pronounced gene body chromatin accessibility in the clusters that also show the highest gene body chromatin accessibility for *ZmWUS1* (Figure 1D). WT C6 and C11, and *Bif3* C8 and C9 were annotated as epidermis, as they exhibited significant gene body chromatin accessibility for *OUTER CELL LAYER4 (OCL4)* ^28^, a transcription factor gene expressed in the meristem L1 layer and in the epidermis (Figure 1E). WT C3-2 and C3-4, along with *Bif3* C1-1 and C1-3, were annotated as suppressed bract or glume primordia, due to increased gene body chromatin accessibility of *ZEA YABBY15* (*ZYB15*) ^8,29^ (Figure 1F). WT C10 and *Bif3* C1 and C2 were classified as the base of the axillary meristem, as they were characterized by notable gene body chromatin accessibility for *RAMOSA1 (RA1)* ^30^ (Figure 1G). The remaining clusters were annotated using previously published cell-type-specific markers (Figure S3). All clusters were identified in both WT and *Bif3*, except for a few unknown clusters that we were unable to annotate and nuclei at different stages of the cell cycle (Figure 2A; Figure S1F-J; Figure S2 C and D).

**Figure 1.**
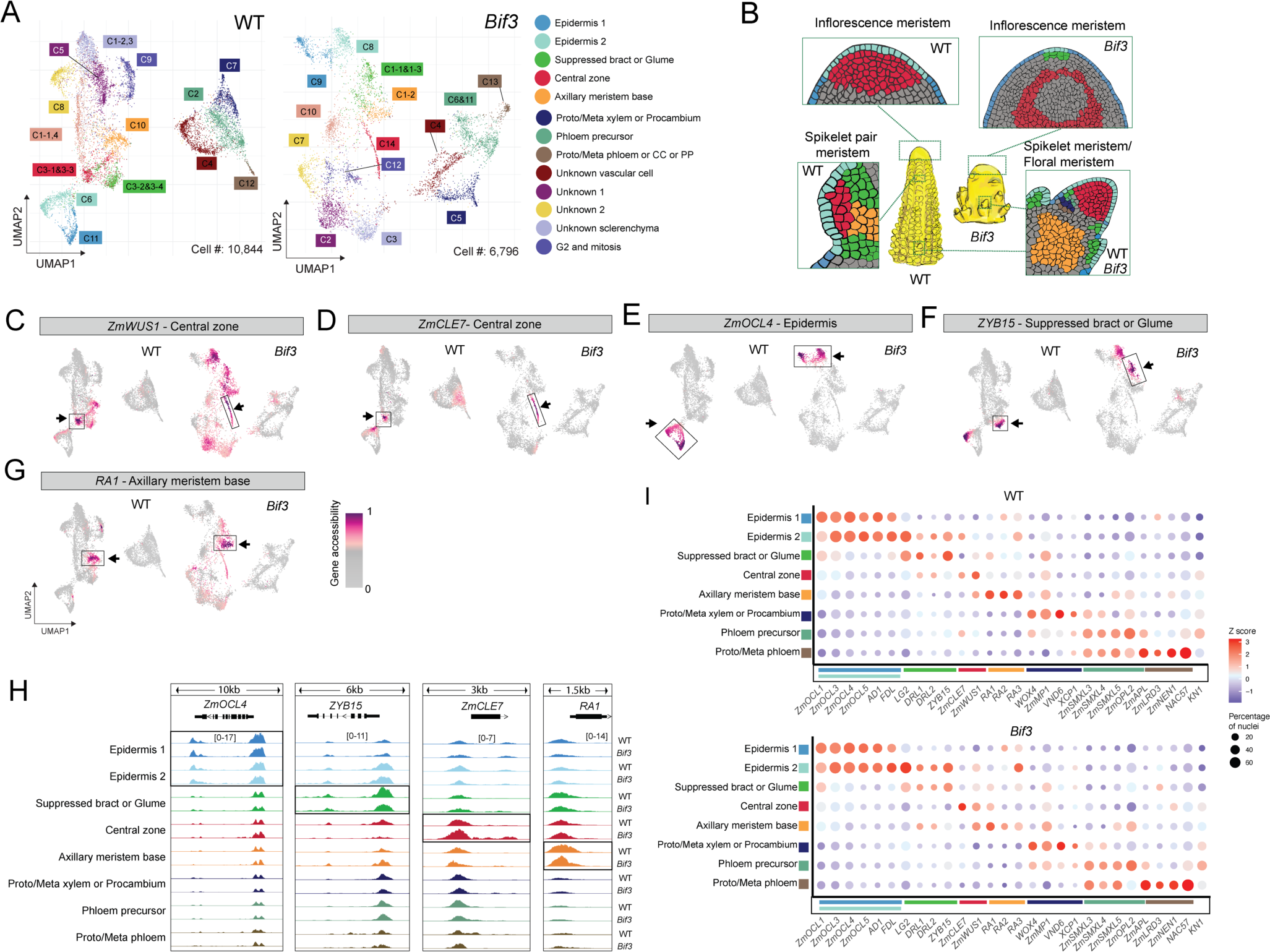
Annotation of central zone nuclei with chromatin accessibility in WT and *Bif3*. **(A)** UMAP plot for WT and *Bif3*, colored to represent the 14 cell types. **(B)** Illustrations depicting the phenotypes of WT and *Bif3*, including longitudinal sections of the inflorescence meristem, the spikelet pair meristem, the spikelet meristem, and the floral meristem are colored to correspond with the cell types in the UMAP. Grey colors represent cell types with an unknown annotation. **(C-G)** UMAP plots highlighting nuclei exhibiting high gene body chromatin accessibility, depicted in purple, surrounding the marker genes of *ZmWUS1* **(C)**, *ZmCLE7* **(D)**, *ZmOCL4* **(E)**, *ZYB15* **(F)**, and *RA1* **(G)**. **(H)** Genome browser track displaying chromatin accessibility around marker genes for meristem-associated cells in WT and *Bif3*. The ranges of number of indicate CPM (counts per million) normalized Tn5 insertion number. **(I)** Gene body chromatin accessibility patterns of 30 marker genes in WT and *Bif3*. The dot size represents the percentage of nuclei that have chromatin accessibility around the marker genes. The values are Z-score normalized aggregated number of Tn5 insertions by cell types.

**Figure 2.**
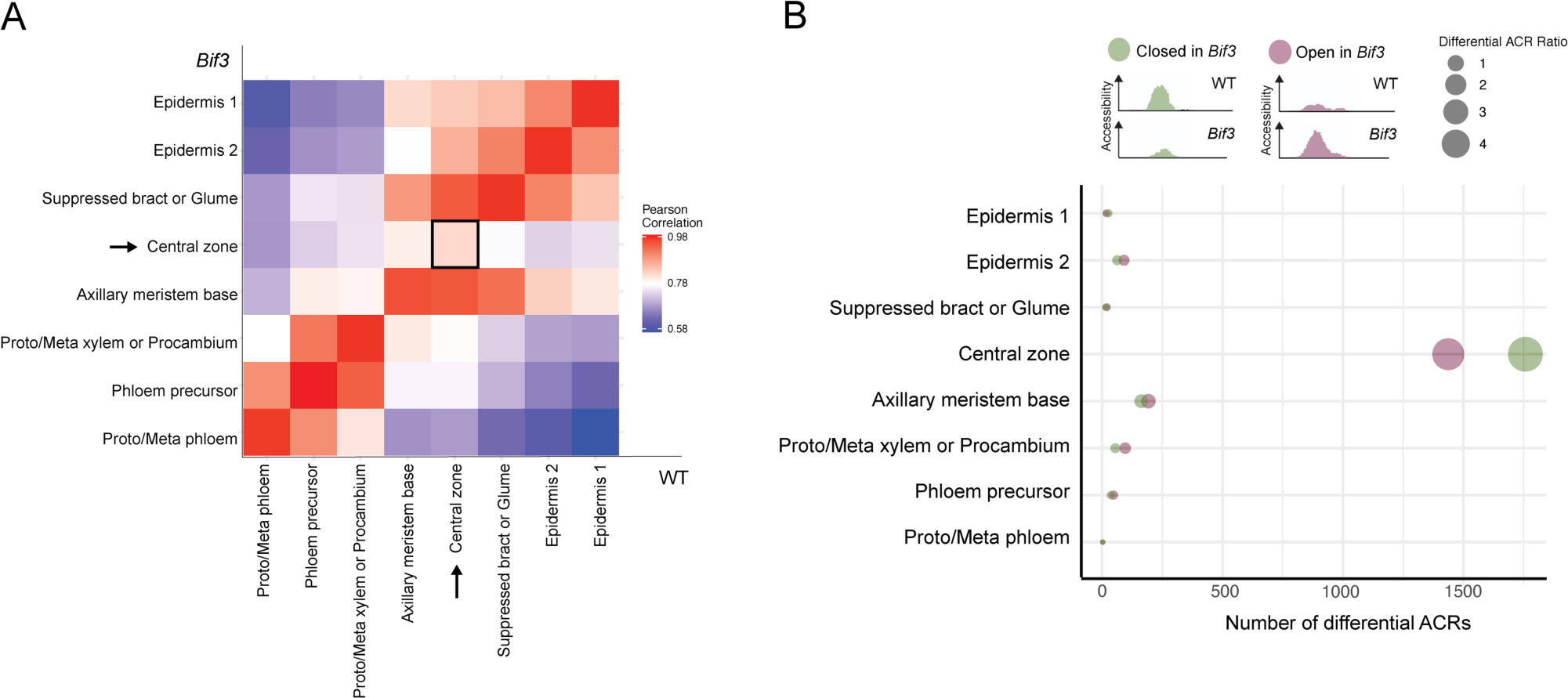
The overall chromatin accessibility profile of WT and *Bif3*. **(A)** A heatmap depicts the correlation of intergenic ACRs between WT and *Bif3*. The 2,000 most variable intergenic ACRs were selected for analysis. The correlation is based on the aggregated number of Tn5 insertions by cell type between WT and *Bif3*. **(B)** A dot plot displays the number of differential ACRs across various cell types. The illustration provides examples of differential ACRs with either higher chromatin accessibility in WT or *Bif3*. The total number of differential ACRs, as well as those exhibiting higher chromatin accessibility in either *Bif3* or WT, are distinctly colored. The ratio of differential ACRs is calculated by dividing the number of differential ACRs by the total number of intergenic ACRs.

Although there are significant morphological differences between WT and *Bif3* ^8^, there was an unexpected amount of consistency in cell-type-specific patterns of chromatin accessibility (Figure 1H; Figure S3). To evaluate the gene body chromatin accessibility pattern more comprehensively for all marker genes, we generated dot plots for 30 marker genes in WT and *Bif3* across all clusters (Figure 1I). The results revealed similar patterns of marker gene chromatin accessibility between WT and *Bif3*, suggesting that the annotated cell types correspond between the two genotypes. Next, using the defined clusters, we identified cell-type-specific markers *de novo* and again we observed a clear correspondence in gene body chromatin accessibility between WT and *Bif3* (Figure S4). These findings indicate that there is a similarity in the patterns of gene body chromatin accessibility for primary marker genes between WT and *Bif3*, which aids in annotating corresponding cell types. The distribution of the proportion of annotated cells aligns with the phenotypic characteristics observed in WT and *Bif3* (Figure S5), where the *Bif3* mutant exhibited a reduced number of cells in the axillary meristem base, coupled with an increased number in the suppressed bract cells ^26^ (Figure 1B).

### Intergenic chromatin accessibility specifically differs in the central zone of *Bif3*

To understand the molecular differences in Accessible Chromatin Region (ACR) that underlie the morphological variation between WT and *Bif3*, we identified differential ACRs, dynamic regions of chromatin accessibility that differ between WT and *Bif3*. Upon merging the ACRs to form a union using ACRs identified from each cell type, the total number of ACRs amounted to 91,386 in WT and 77,393 in *Bif3* (Table S4). In WT, approximately 46.6% of the ACRs are gene-overlapping ACRs, 36.6% are distal ACRs – located more than 2 kbp from the closest gene, and 16.7% are proximal ACRs – situated less than 2 kbp from a gene. Similarly, in *Bif3*, the distribution includes ∼47.3% gene-overlapping ACRs, ∼35.6% distal ACRs, and 17% proximal ACRs.

We investigated genome-wide accessible chromatin changes within cell types in WT and *Bif3* (Figure 2A). We created intergenic ACR sets, defined by the condition that the midpoint of the ACR does not overlap with any genic region. We aggregated intergenic ACRs across all cell types and genotypes, resulting in 58,578 intergenic ACRs, each spanning 500 bp based on the peak summit. Correlations of read density within ACRs between corresponding cell types exceeded 0.95 for all the cell types except the central zone, indicating a high degree of similarity between WT and *Bif3*. The correlations between cell types are much higher than those between unrelated cell types. The only cell type that did not follow this trend in WT and *Bif3* was the central zone nuclei, which displayed a correlation of 0.81. This suggests drastic alterations in chromatin accessibility within the *Bif3* central zone, amidst an otherwise conserved chromatin landscape, suggesting the central zone is the source of the morphological differences observed in *Bif3*.

To further investigate whether chromatin accessibility in the central zone of *Bif3* is increased or decreased, we identified differentially accessible chromatin regions (differential ACRs). For each cell type, we combined intergenic ACRs from both WT and *Bif3*, covering the full extent of ACRs found in either genotype. The procedure was performed independently for each cell type, resulting in intergenic ACR sets of varying numbers. On average, 40,208 intergenic ACRs were pairwise tested for differential ACRs by comparing their chromatin accessibility between WT and *Bif3*. Most cell types had fewer than 355 differential ACRs, representing less than 1% of the total ACRs per cell type, whereas the central zone exhibited 3,194 differential ACRs, accounting for 8.5% of its total ACRs (Figure 2B). Among the differential ACRs in the central zone, 1,437 showed increased chromatin accessibility in *Bif3* (termed ‘increased in *Bif3*’), whereas 1,757 exhibited decreased chromatin accessibility in *Bif3* (termed ‘decreased in *Bif3*’) (Table S5).

### Regions with increased chromatin accessibility due to overexpressed *ZmWUS1* were enriched for the CAATAATGC motif and genes near this motif show signs of elevated transcription

To uncover transcription factors potentially involved in the altered chromatin accessibility landscape, we identified enriched motifs in differential ACRs increased and decreased in *Bif3*, respectively. The differential ACRs increased in *Bif3* were enriched for a single significant motif (CAATAATGC), whereas the differential ACRs decreased in *Bif3* showed enrichment for five distinct motifs (Figure 3A). In total, there were 381 differential ACRs (∼26.51% of those increased in *Bif3*) that had a CAATAATGC motif. The CAATAATGC motif has a core motif of TAAT, which was previously identified as a WUS binding motif ^13,18^. This finding is supported by our hypothesis that overexpressed ZmWUS1 targets chromatin regions not originally intended as its binding sites in WT, thereby increasing chromatin accessibility. Thus, the ‘increased in *Bif3*’ differential ACRs containing the CAATAATGC motif in the central zone represent ACRs directly associated with the overexpression of *ZmWUS1*.

**Figure 3.**
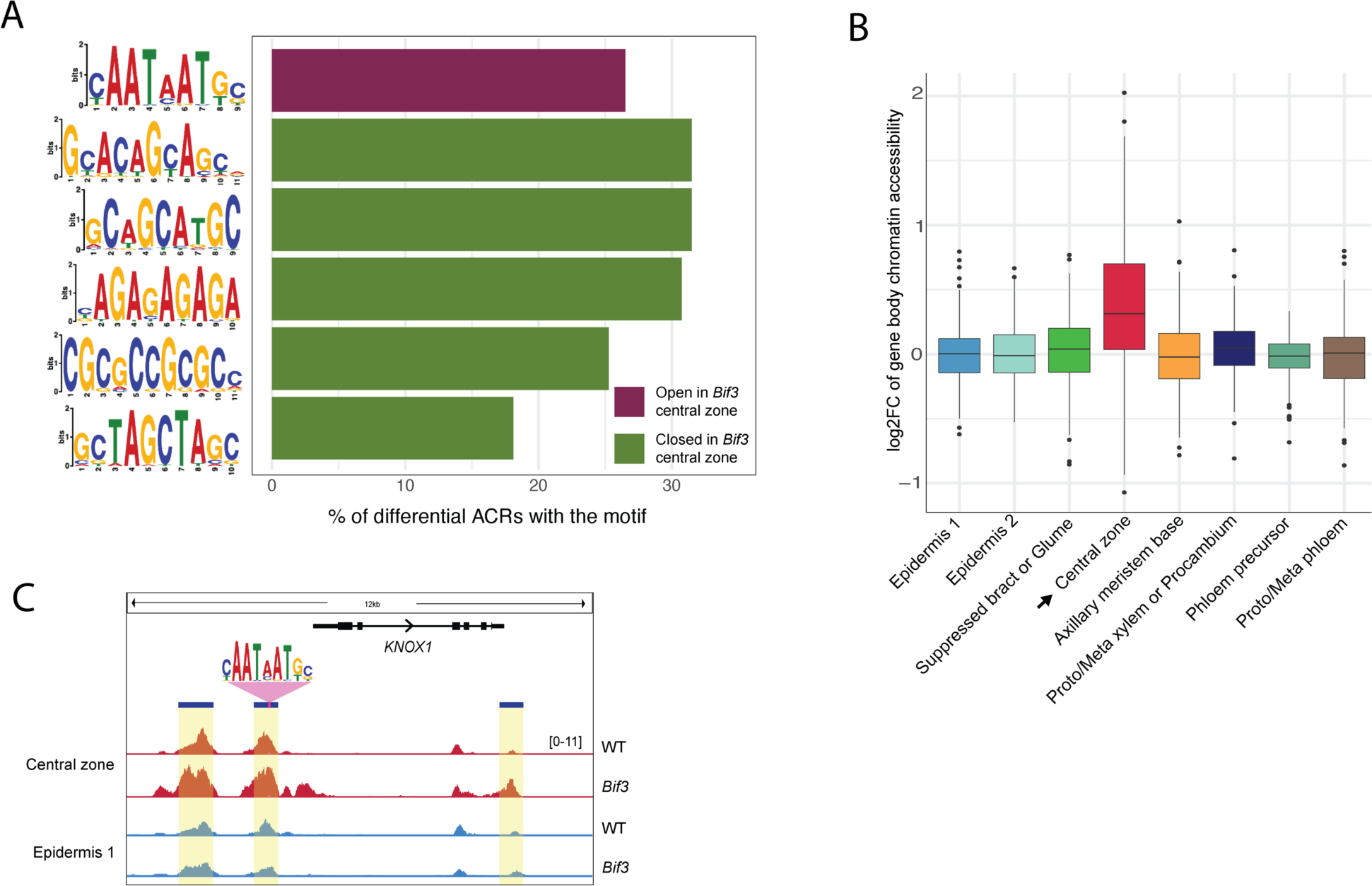
Characteristics of differentially Accessible Chromatin Regions (differential ACRs) between WT and *Bif3* by cell types. **(A)** The PWM illustrates significant motifs discovered within differential ACRs in the central zone (E-value <1). The ratio indicates the number of differential ACRs containing the motif divided by the total number of differential ACRs. The color denotes whether the differential ACR sets are increased or decreased in *Bif3* central zone. **(B)** Box plot displays the log2 fold change (log2FC) of gene body chromatin accessibility surrounding differential ACRs increased in *Bif3* with the CAATAATGC motif. Log2FC was calculated by normalizing gene body chromatin accessibility of differential ACRs in *Bif3* to those of WT. Rows represent individual genes, while columns represent cell types. **(C)** Genome browser view illustrates chromatin accessibility in central zone for the peaks with the CAATAATGC motif.

*ZmWUS1* is a member of the *WUSCHEL-related homeobox* (*WOX*) gene family that shares a conserved DNA-binding homeodomain ^31^. For example, *in vitro* assays revealed interactions between TAAT sequences and OsWOX5 (*Oryza sativa*) in rice ^32^, and TTAATT and Arabidopsis WOX11 ^24^. One possibility for the detection of the TAAT motif in the differential ACRs is that ZmWUS1 enhances the expression of other *WOX* genes, making it difficult to distinguish if these ACRS are affected by ZmWUS1 or WOX transcription factors. Thus, we explored which cell types have higher gene body chromatin accessibility, a proxy for expression ^33^, in the 14 annotated *WOX* genes ^34^ and *ZmWUS2* ^35^, the *ZmWUS1* co-ortholog. Our analysis in WT showed that although most *WOX* genes have increased chromatin accessibility in vascular nuclei, *ZmWOX2A, ZmWOX9B, ZmWOX9C, ZmWOX13B*, and *ZmWUS2* showed greater chromatin accessibility in the central zone nuclei compared to other cell types (Figure S6A). A comparison of gene body chromatin accessibility between WT and *Bif3* showed a general decrease of chromatin accessibility for *WOX* genes in the *Bif3* central zone, particularly *ZmWOX3A*, *ZmWOX9B,* and *ZmWOX9C* (Figure S6B and C). This is highlighted by the ACR near the promoter of *ZmWUS2*, which exhibits a substantial reduction in chromatin accessibility in the *Bif3* central zone nuclei (Figure S6D) and is downregulated in *Bif3* RNA-seq datasets ^8^. These findings suggest that the prevalence of CAATAATGC motifs in the ‘increased in *Bif3*’ differential ACRs is not predominantly driven by other *WOX* family members.

We investigated whether the 381 differential ACRs in the central zone nuclei with a CAATAATGC motif are associated with activated or inactivated nearby genes. We used gene body chromatin accessibility proximal to these differential ACRs targets as proxy for whether a gene is expressed or not, as previous research showed there is a good correlation ^33^. We discovered that gene body chromatin accessibility of most of the genes associated with the 381 differential ACRs increased in the central zone of *Bif3* compared to WT (Figure 3B). The central zone displayed an average log2 fold change in gene body chromatin accessibility, comparing *Bif3* to WT, of approximately 0.38. In contrast, other cell types displayed log2 fold change values that, on average, were less than 0.05 in magnitude, indicating minimal changes. As examples, we observed a differential ACRs that becomes accessible in *Bif3* in the central zone near *KNOTTED-like homeobox* (*KNOX1)*, a meristematic gene ^24^ (Figure 3C). This indicates that the differential ACRs exhibiting increased chromatin accessibility in *Bif3* are associated with elevated transcription of nearby genes.

### Accessible chromatin regions with decreased accessibility due to overexpressed *ZmWUS1* are associated with reduced auxin signaling

We examined gene body chromatin accessibility changes near differential ACRs that were decreased in *Bif3* in the central zone nuclei, which were associated with five different motifs: GCACAGCAGC, GCAGCATGC, CGCGCCGCGCC, GCTAGCTAGC, and AG repeats (Fig 3A). The multiple motifs present in the decreased ACR set suggests that distinct transcription factors may be involved with these regions, each recognizing their specific motif sequence. Near these decreased differential ACRs with the five motifs, we observed decreased gene body chromatin accessibility in *Bif3* central zone genes (Figure S7). This indicates that differential ACRs decreased in *Bif3* are likely involved in reducing the transcription of nearby genes.

Gene ontology analysis revealed that the differential ACRs decreased in *Bif3* in the central zone were linked to transcription regulation, developmental regulation and auxin hormone responses (Figure 4A). As meristem size regulation is associated with auxin hormone pathways ^36^, we investigated *AUXIN RESPONSE FACTORS* (*ARFs*) genes. We discovered 10 *ZmARF* genes nearby differential ACRs in the central zone. All these differential ACRs decreased in *Bif3*, whereas none were nearby ACRs increased in *Bif3* (Figure 4B; Figure S8A). Most of these *ZmARF* genes displayed decreased gene body chromatin accessibility in the *Bif3* central zone nuclei, with varying changes observed in other cell types. Genome browser views showed reduced chromatin accessibility of differential ACRs upstream of *ZmARF4* and *ZmARF23*, as well as reduced gene body chromatin accessibility in the *Bif3* central zone (Figure 4C). Thus, differential ACRs decreased in central zone of *Bif3* are associated with likely decreased transcription of *ZmARFs*. This indicates that *ZmARFs* transcription is negatively associated with overexpressed *ZmWUS1* and is consistent with the negative regulation of *ZmARFs* being associated with increased meristem size ^37^.

**Figure 4.**
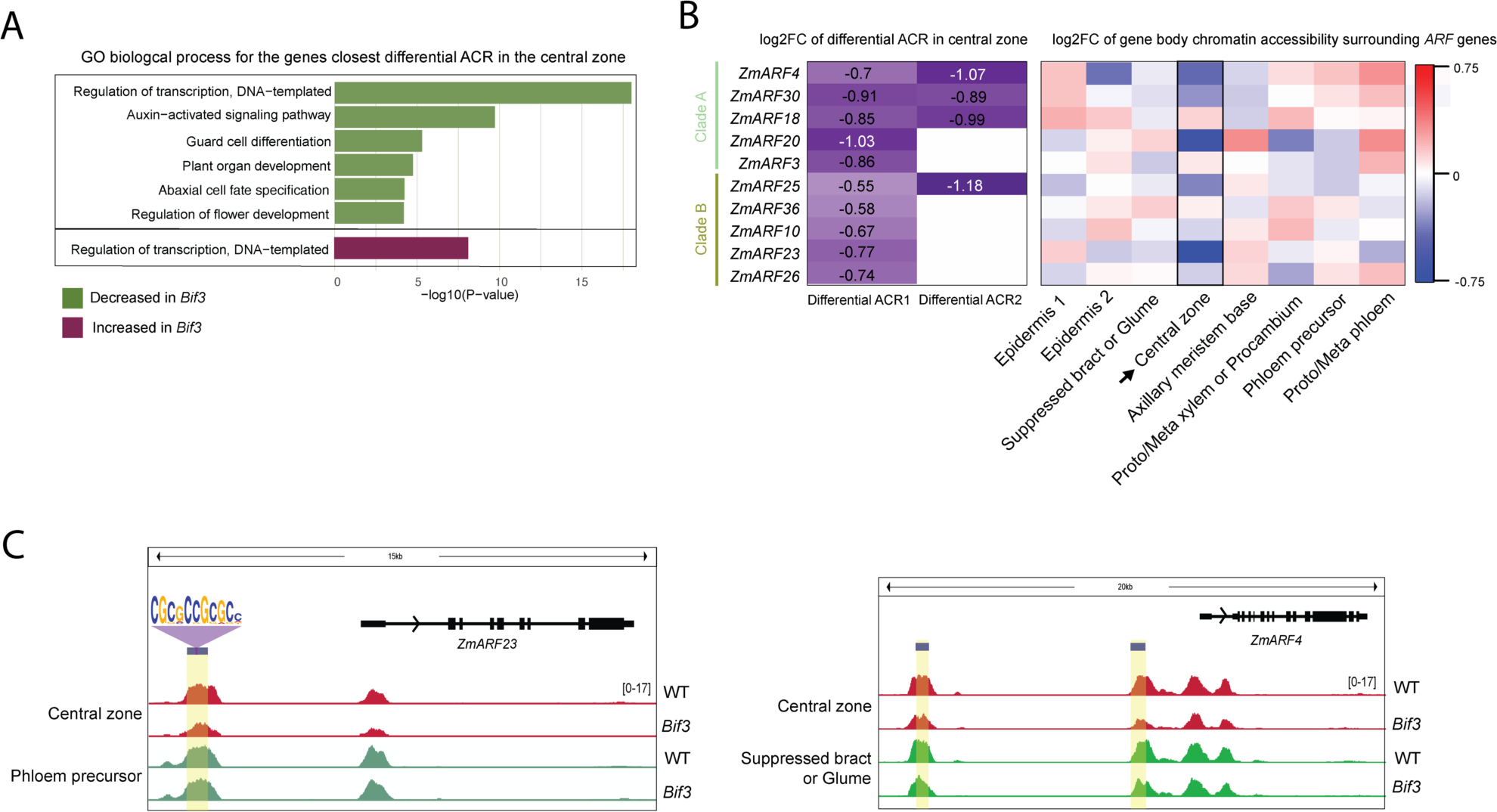
The *ARF* genes are associated with decreased differential ACRs by overexpression of *ZmWUS1*. **(A)** Bar graph illustrates significant biological process terms identified using a Fisher’s exact test using the genes closest to the differential ACRs in the central zone (FDR <0.05). **(B)** The left heatmap shows the logFC of differential ACRs around *ARF* genes, whereas the right heatmap shows the gene body chromatin accessibility of *ARFs* by cell types. **(C)** Chromatin accessibility in the central zone around *ZmARF4* and *ZmARF23*. The blue bar and yellow highlighted region indicate the differential ACRs.

We further categorized *ZmARFs* near differential ACRs in the central zone into clade A and B; clade A, are transcriptional activators, while clade B are repressors ^38,39^. Among the 10 *ZmARFs* found near differential ACRs, *ZmARF4*, *ZmARF18*, *ZmARF20*, and *ZmARF30* belong to clade A, whereas *ZmARF10*, *ZmARF23*, *ZmARF25*, and *ZmARF36* are in clade B ^40^. The differential ACRs near clade A *ZmARFs* lacked the five motifs found in the decreased differential ACRs, whereas those near clade B *ZmARFs* contained at least one of these motifs. Notably, all differential ACRs near clade B *ZmARFs* commonly featured the CGCGCCGCGCC motif (Figure S8B). This indicates that the negative regulation of *ZmARFs* by overexpressed *ZmWUS1* involves both clade A and clade B, with clade B being predominantly regulated by a single CGCGCCGCGCC motif.

### ZmWUS1 DAP-seq identified the TGAATGAA motif rather than the CAATAATGC motif

We used DAP-seq ^24^ to identify DNA sequences directly bound by ZmWUS1 across the maize genome (Figure 5A). In total, 10,382 peaks were identified, which uncovered TGAATGAA as the top-ranked motif consistently observed at the center of the peaks (Figure 5B). The first occurrence of TGAA in the motif shows strong representation, whereas the second occurrence exhibits variability especially in the tail of the motif (Figure 5C). The analysis using Tomtom ^41^ and the JASPAR motif database for plants ^42^ and the DAP-seq cistrome database ^24^ revealed that the TGAATGAA motif from ZmWUS1 DAP-seq shares similarity with the Arabidopsis WUS motif (E-value=0.079), but not with any other motif. The TGAATGAA motif matches the motif previously identified using ChIP-seq and DAP-seq using Arabidopsis WUS ^8,24,25^. This shows the TGAATGAA motif is unique from other known TF motifs in plants.

**Figure 5.**
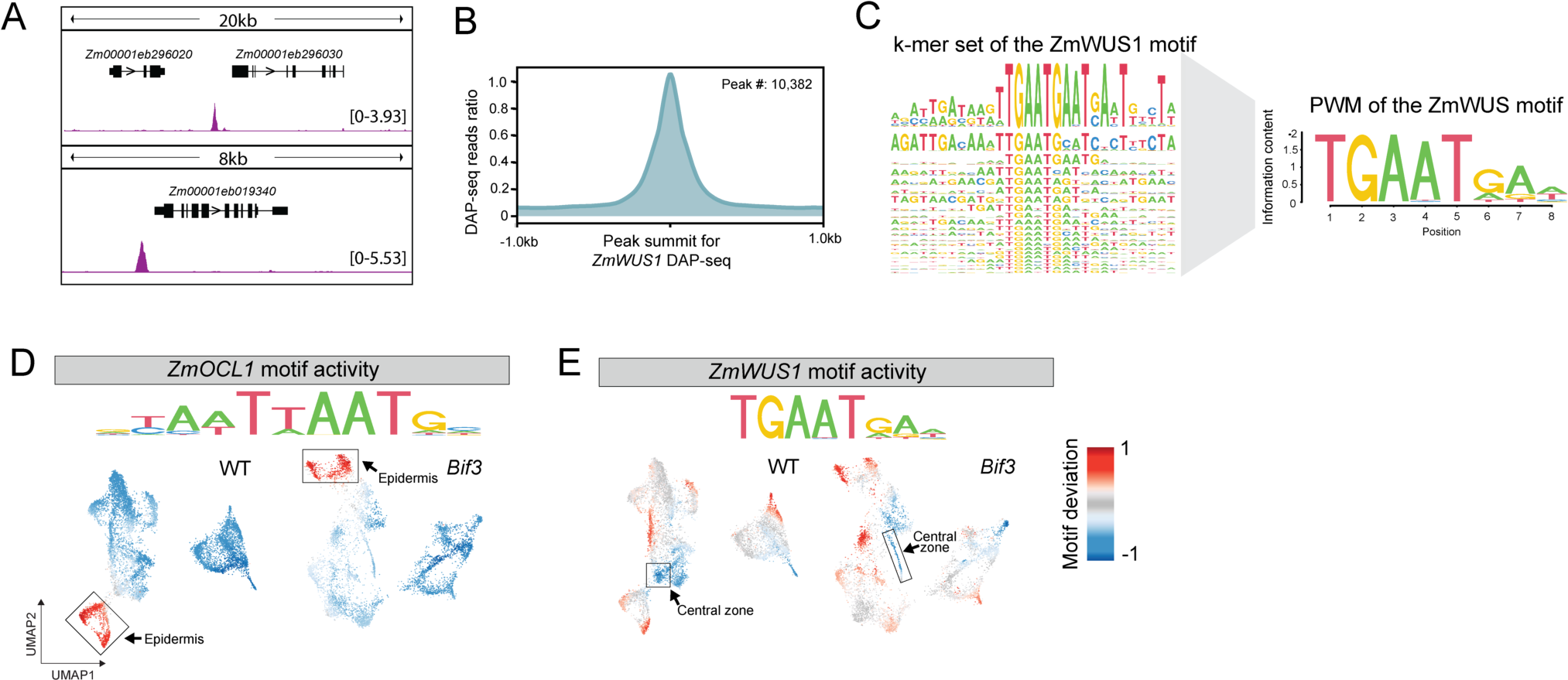
The ZmWUS1 motif identified using DAP-seq shows different activity within ACRs depending on the cell type. **(A)** A genome browser view showing the peaks from ZmWUS1 DAP-seq. The y-axis shows CPM values. **(B)** A metaplot displaying the ratio of reads within a 1-kilobase pair region surrounding the ZmWUS1 DAP-seq peaks. **(C)** Identification of the ZmWUS1 motif using a k-mer set analysis, along with Position Weight Matrix (PWM) models of the most enriched and significant motifs. **(D)** The motif deviation of ZmOCL1 in WT and *Bif3*. The motif is known for Arabidopsis HDG1, which is orthologous to ZmOCL1. The box with arrows indicates the epidermis cells. **(E)** The motif deviation of ZmWUS1 in WT and *Bif3*. The box with arrows indicates the central zone cells.

We hypothesized that the TGAATGAA motif would be accessible for binding in the nuclei where *ZmWUS1* is active. First, we identified all intergenic ACRs across all cell types in each genotype, which revealed 58,424 and 49,618 ACRs in WT and *Bif3* respectively. Next, we used chromVar ^43^ to calculate motif deviation scores, which represents the likelihood of observing a specific motif with high chromatin accessibility within the intergenic ACRs in each nucleus. Our previous study indicated that in certain cell types, transcription factors with high gene body chromatin accessibility also tend to have higher motif deviation scores ^33^. As an example, based on its orthology to Arabidopsis HOMEODOMAIN GLABROUS1 (HDG1), the *ZmOCL1* transcription factor is expressed in the outer layers and is predicted to bind a TAATTAATG motif. Motif deviation for *ZmOCL1* indicated its target motif is enriched in the intergenic ACRs of the epidermis in both WT and *Bif3*, as its motif deviation is higher in epidermis compared to other cells (Figure 5D). This suggests that the ZmOCL1 transcription factor, expressed in the epidermis, binds to ACRs within the same cells where it is expressed.

For the TGAATGAA motif identified via DAP-seq for ZmWUS1, we observed low motif deviation scores in the central zone of both WT and *Bif3* (Figure 5E). This indicates that TGAATGAA motifs are less prevalent in the accessible chromatin regions (ACRs) of the central zone where *ZmWUS1* is expressed, compared to other cell types. In both genotypes, the highest motif deviation scores for ZmWUS1’s TGAATGAA motif was in the epidermis, proto/meta xylem or procambial cells. This suggests that ZmWUS1 likely targets cis-regulatory elements with the TGAATGAA motif in specific cell types away from the central zone if it can act as a monomer or homodimer.

## Discussion

ZmWUS1 plays a crucial role in meristem development as a transcription factor that binds to cis-regulatory elements and facilitates formation of the ear. *ZmWUS1* expression is confined to the central zone, which has presented challenges in identifying associated *cis*-regulatory elements. To address this, we compared cell-type chromatin accessibility differences in the central zone between WT and *Bif3*, identifying 1,437 regions with increased chromatin accessibility and 1,757 regions with decreased chromatin accessibility in *Bif3*. The accessible chromatin regions with increased accessibility in the *Bif3* central zone nuclei were enriched for the CAATAATGC motif, comprising approximately 26.51% of these regions. WUS can bind TAAT repeats in a variety of *in vitro* binding assays ^14,20^. Our data suggests that overexpressed *ZmWUS1* might lead to binding *cis*-regulatory elements in accessible chromatin beyond its typical targets when expressed at WT levels. Together, these results suggest that ZmWUS1 likely targets TAAT in the central zone nuclei. Another possibility is that related gene family members that also target the TAAT motif perform this function; however, the diminished gene body chromatin accessibility observed for *ZmWUS2* or *WOX* genes reduces this likelihood and supports the hypothesis that ZmWUS1 targets the CAATAATGC motif in the central zone nuclei (Figure S6). As WUS is proposed to be a dual functional transcription factor that can activate or repress gene expression in Arabidopsis depending on its context^16^, we examined gene body chromatin accessibility changes near target regions harboring the CAATAATGC motif and discovered that their gene body chromatin accessibility mostly increases, which implies that ZmWUS1 likely acts as an activator in the central zone nuclei in maize.

The TGAATGAA motifs identified by ZmWUS1 DAP-seq were not predominant in the accessible chromatin regions of *Bif3* in the central zone nuclei; instead, the CAATAATGC motif was prevalent. DAP-seq captures ZmWUS1 binding, either as a monomer or a homodimer ^24^, and suggests that the TGAATGAA motif might derive from either form of ZmWUS1 binding. Protein structural analysis indicates a potential for homodimer formation of ZmWUS1 using TGAATGAA for DNA binding ^8^. Given the increased amount of ZmWUS1 in the central zone nuclei of *Bif3*, there’s a heightened likelihood for homodimer formation. Absence of the TGAATGAA motifs in accessible chromatin regions in the central zone nuclei in *Bif3* suggests additional factors influence ZmWUS1 binding to target sequences in the central zone. One possibility is that there could be protein interactions with the other transcription factors that are specific to the central zone and restrict the binding of ZmWUS1 in comparison to nearby cells where ZmWUS1 is able to act as a monomer or homodimer (Figure 6). Alternatively, cell-type-specific post-translational modifications of ZmWUS1 could affect its activity including its ability to partner with other proteins or it could affect its DNA-binding affinity ^44,45^. Future research that incorporates cell-type-specific binding of ZmWUS1 to its targets will be needed to resolve this question.

**Figure 6.**
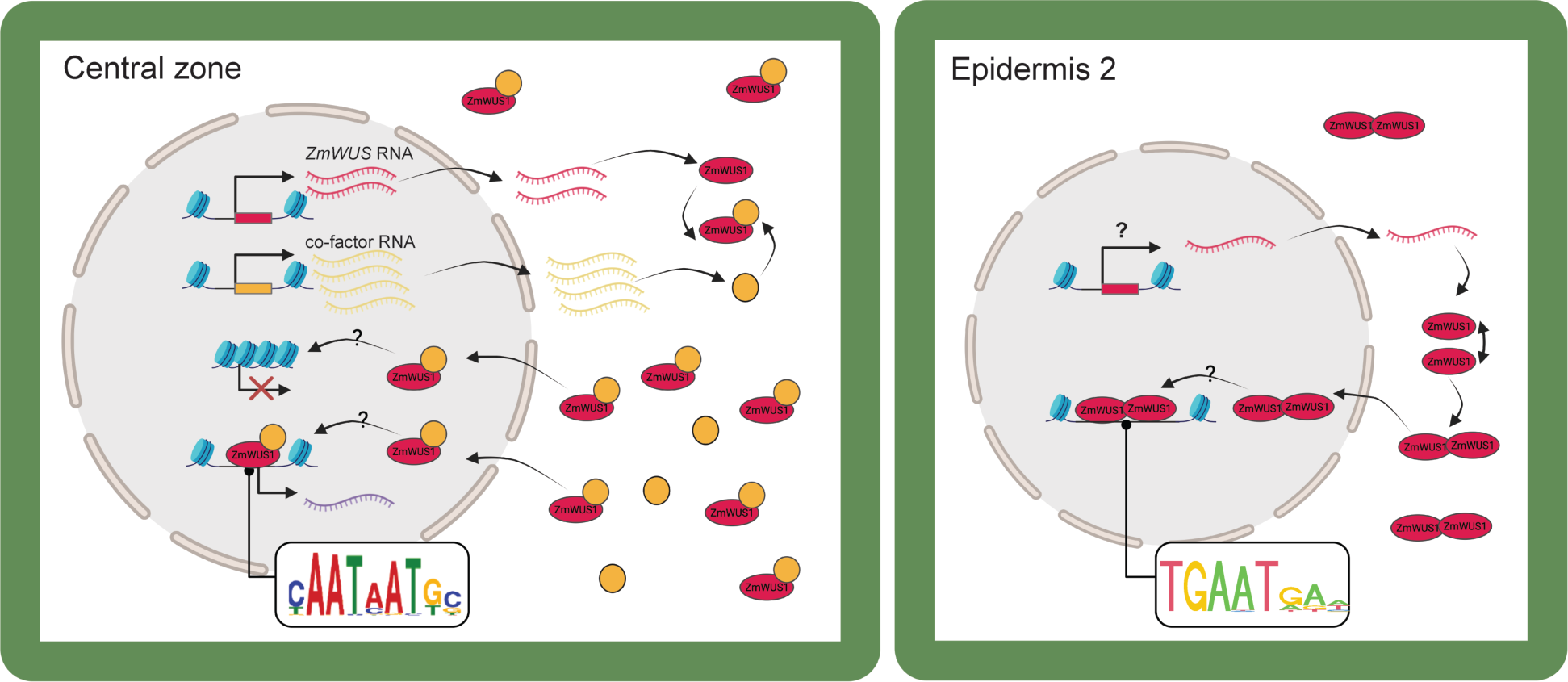
A proposed model for central zone specific ZmWUS1 *cis*-regulation. cell-type-specific motif usage by ZmWUS1 could be attributed to the unique molecular interactions occurring in central zone cells.

The TGAATGAA motif was found more frequently in the ACRs of the epidermis and proto/meta xylem cells compared to those of central zone in both WT and *Bif3.* This includes the ‘epidermis 2’ nuclei, which had gene body chromatin accessibility for *ZmWUS1*. This observation suggests that *ZmWUS1*, when expressed in epidermis 2, may target the TGAATGAA motif. However, the enriched TGAATGAA motif in ACRs of epidermis 1 and the proto/meta xylem or procambium cells remains challenging to explain. In the absence of *ZmWUS1* expression in these cells, one might speculate that ZmWUS1 diffuses there. Nonetheless, even if ZmWUS1 does move, its effects seem minimal. Furthermore, the coinciding expression patterns of *ZmWUS1* and *ZmCLE7* within the central zone of maize immature ears prompt further inquiry into the necessity and scope of ZmWUS1’s movement for different motif usage. Our findings lead to a proposed model illustrating cell-type-specific motif utilization by ZmWUS1 (Figure 6).

The decreased chromatin accessibility in the central zone nuclei due to overexpressed *ZmWUS1* raises intriguing questions. If ZmWUS1 actively binds these regions, they should be accessible, yet they are inaccessible, indicating a past binding event or possibly indicating that the regions were never accessible in the first place. One possibility is that overexpressed *ZmWUS1* previously initiated chromatin closure, an event that might have occurred before sampling of the *Bif3* tissue used in this study. This is plausible given *ZmWUS1* is expressed during the embryonic stage ^35^. Given that the central zone encompasses multiple cell types, ZmWUS1 could potentially interact with other transcription factors across these different cells. This interaction is suggested by the enrichment of five distinct motifs (GCACAGCAGC, GCAGCATGC, CGCGCCGCGCC, GCTAGCTAGC, and AG repeats) in the decreased chromatin accessibility region in the central zone. Alternatively, ZmWUS1 might indirectly cause chromatin closure without directly binding to these regions. For example, the presence of AG repeats is one class of polycomb response elements ^46,47^, and suggests a potential silencing role directed by the Polycomb repressive complex 2. The decreased gene body chromatin accessibility of genes linked to these motifs supports a silencing function, pointing to a complex interplay where ZmWUS1 might be implicated in silencing either directly or indirectly. Future studies that can dissect cell-type-specific binding of ZmWUS1 will be needed to resolve these possibilities.

The reduced chromatin accessibility near *ZmARF* genes suggests a link between overexpressed *ZmWUS1* and diminished auxin signaling in the central zone nuclei. This observation aligns with the known interactions between *Bif3* and auxin-insensitive mutants^8^. In Arabidopsis, the shoot apical meristem exhibits lower auxin signaling in the central zone compared to peripheral regions ^48^. Previous research indicates that induced *WUS* expression further decreases auxin signaling in the central zone, suggesting WUS shields stem cells from auxin-driven differentiation ^25^. This auxin-related regulatory mechanism implies that ZmWUS1 might negatively influence *ZmARF* gene regulation, affecting auxin signaling pathways. The diminished transcription of *ZmARF* in the central zone accounts for the reduced differentiation observed in *Bif3*, characterized by an increased stem cell count in the inflorescence meristem and the presence of single rather than paired spikelet meristems. Overall, this study highlights how the overexpression of *ZmWUS1* influences early ear development by enhancing chromatin accessibility in central zone ACRs containing the CAATAATGC motif and simultaneously reducing ACRs associated with the auxin signaling pathway.

## STAR*METHODS

### RESOUCE AVAILABILITY

#### Lead contact

Further information and requests for resources and reagents should be directed to and will be fulfilled by Lead Contact, Robert J. Schmitz.

#### Materials availability

This study did not generate new unique reagents.

#### Data and code availability

Single-cell ATAC-seq and DAP-seq data have been deposited at the NCBI GEO database under accession code GSE266007.

### EXPERIMENTAL MODEL AND SUBJECT DETAILS

Immature ears (2-6 mm in size) were collected from the inbred line A619 wild-type, and +/*Bif3* plants (back-crossed 12 times in A619) grown in winter 2020 in the Waksman Institute greenhouse (Piscataway, NJ, USA), and in summer 2021 in the Waksman Institute field (Piscataway, NJ, USA). The *Bif3* mutant phenotype is completely dominant in A619 ears ^26^. Immature ears were harvested (36 and 31 samples for A619, and 41 and 36 for +/Bif3 in 2020 and 2021, respectively) approximately at the same time of the day (from 11am-2pm) and bulked for subsequent analysis. Greenhouse sown seeds were grown in 5-gallon pots filled with PROMIX BX General Purpose Soil and supplemented with 14-14-14 Osmocote. Soil was saturated with tap water and placed under GE Lucalox 400W lighting. Seedlings were grown under a photoperiod of 16 hours of light, 8 hours of dark. The temperature was maintained at ∼25°C during light hours with a relative humidity of approximately 30%. Greenhouse growing conditions were monitored and controlled by MicroGrow Greenhouse Systems.

### METHOD DETAILS

#### Single-cell ATAC-seq library construction

The protocol for nuclei isolation and purification was adapted from a previously described method with specific modifications to improve nuclei yield and quality ^49^. In brief, approximately 5-6 immature maize ears were fine chopped on ice for approximately one minute using 300 μL of pre-chilled Nuclei Isolation Buffer (NIB, 10 mM MES-KOH at pH 5.4, 10 mM NaCl, 250 mM sucrose, 0.1 mM spermine, 0.5 mM spermidine, 1 mM DTT, 1% BSA, and 0.5% TritonX-100). Following chopping, the mixture was filtered through a 40-μm cell strainer and then centrifuged at 500 rcf for 5 minutes at 4°C. The supernatant was carefully removed, and the pellet was washed in 500 μL of NIB wash buffer, comprised of 10 mM MES-KOH at pH 5.4, 10 mM NaCl, 250 mM sucrose, 0.1 mM spermine, 0.5 mM spermidine, 1 mM DTT, and 1% BSA. Then, the sample was filtered again through a 20-μm cell strainer, and then gently layered onto the surface of 500 μL of a 35% Percoll buffer, prepared by mixing 35% Percoll with 65% NIB wash buffer, in a 1.5-mL centrifuge tube. The nuclei were centrifuged at 500 rcf for 8 minutes at 4°C. After centrifugation, the supernatant was carefully removed, and the pellets were resuspended in 10 μL of diluted nuclei buffer (DNB, 10X Genomics Cat# 2000207). Approximately 5 μL of nuclei were diluted 10 times, stained with DAPI (Sigma Cat. D9542), and then assessed for quality and density with a hemocytometer under a microscope. The original nuclei were diluted with DNB buffer to a final concentration of 3,200 nuclei per uL. Finally, 5 uL of nuclei (16,000 nuclei in total) were used as input for scATAC-seq library preparation. scATAC-seq libraries were generated using the Chromium scATAC v1.1 (Next GEM) kit from 10X Genomics (Cat# 1000175), following the manufacturer’s instructions (10xGenomics, CG000209_Chromium_NextGEM_SingleCell_ATAC_ReagentKits_v1.1_UserGuide_RevE). The libraries were sequenced with Illumina NovaSeq 6000 in dual-index mode with 8 and 16 cycles for i7 and i5 index, respectively.

#### DAP-seq library construction

DAP-seq libraries for assessing ZmWUS1 binding to maize genomic DNA were created as described in Galli et al., 2018 ^40^. In brief, genomic DNA was extracted from the leaves of 14 day-old B73 seedlings using a phenol:cholorform:Iso amyl alcohol procedure. 5ug of extracted DNA was sheared in a Covaris S2 sonicator to 200bp fragments. The fragmented DNA was then end repaired using the END-IT DNA repair kit (Lucigen) per the manufacturer’s recommendations and cleaned using a QiaQuick PCR clean-up method (Qiagen). Eluted DNA was then A-tailed using Klenow exo-(NEB) for 30min at RT. Samples were cleaned using a QiaQuick PCR-clean-up method. Eluted DNA was ligated to truncated Illumina Y-adapters overnight at 16 degrees C using T4 DNA ligase (NEB). Samples were heat inactivated for 10 min at 70 degrees C followed by a 1:1 bead clean-up using AmpureXP beads (Beckman-Coulter). Libraries were quantified using the Qubit HS kit. 1ug of this library was then incubated with in vitro expressed HALO-WUS1 protein bound to 10ul of MagneHALO beads (TNT rabbit reticulocyte expression system and MagneHALO beads, Promega) generated from 1ug of pIX-HALO-WUS1 plasmid. Unbound DNA was washed away using six washes of phosphate buffered saline (PBS), and bound DNA was eluted from the beads using 30ul of 10mM Tris (EB) and heating to 98 degrees C for 10 min. DNA was barcoded and enriched with 19 cycles and Illumina primers prior to sequencing on a NextSeq500 with 75bp Single End reads.

### QUANTIFICATION AND STATISTICAL ANALYSIS

#### High-quality nuclei identification

The reads from scATAC-seq are mapped using the CellRanger-atac count (v2.0.0) by 10X Genomics to modified B73 v5 reference genome. In B73 v5, the scaffolds are removed while Mt (mitochondria) and Pt (plastids) were added from v3 genome. The multi-mapped reads with BWA added tags of “XA” and “SA” were removed ^50^. De-duplication was performed by picard MarkDuplicates (v2.16.0) (https://broadinstitute.github.io/picard/). Single-base pair Tn5 integration were identified using a python script ^51^.

Socrates, R package (https://github.com/plantformatics/Socrates), was used to filter out nuclei that did not meet quality thresholds ^33^. These criteria included the quantification of Tn5 transposase insertions, the proximity ratio of Tn5 insertions to the transcription start site (TSS), the fraction of reads in peaks (FRiP) score, and the organelle DNA ratio. We excluded nuclei falling below the knee point identified in the plot correlating unique Tn5 integration sites per nucleus with barcode rank. Nuclei were filtered if they exhibited TSS ratio or FRiP score beyond three standard deviations from the mean, and if the organelle DNA ratio exceeded 5%. Additionally, nuclei with fewer than 100 accessible chromatin regions, or with an open chromatin peak ratio below 0.01 or above 0.05, were excluded from subsequent analysis.

We used the presence or absence of Tn5 insertions within 500 bp windows as features for dimensionality reduction. A blacklist generated from comparative ATAC-seq to genomic DNA and control ChIP-seq data was applied to remove features. The cleanData function in the Socrates package was used to apply feature filters based on their frequency across the cell population. Features present in more than 5% of cells were retained using the ‘max.t=0.05’ parameter, while those observed in fewer than 1% of cells were excluded with the ‘min.t=0.01’ threshold. Feature normalization was performed using the term frequency-inverse document frequency transformation (TF-IDF) ^52^. The top third of the features, identified as highly variable, were selected for further dimensionality reduction using the reduceDims function in Socrates.

Singular value decomposition (SVD) was used to reduce dimensionality and compute the principal components ^53^. The top 100 principal components were used for Uniform Manifold Approximation Projection (UMAP). Doublets were identified and removed using Scrublet using the software Scrublet as implemented in detectDoublets and filterDoublets function in Socrates with the option of filterRatio=1.5 ^54^. Finally, to integrate the two replicates, we applied the Harmony algorithm with ‘theta=2, sigma=0.1, max.iter.cluster=100, and max.iter.harmony=30’ ^55^. Cell clustering was performed using Louvain clustering on k=50 nearest neighborhood graph with a resolution of 0.3. Subsequent cluster assignment by Louvain community detection identified distinct cell clusters.

#### Gene body chromatin accessibility analysis

Gene body accessibility served as a basis for cell annotation, leveraging the variability across cells. We used marker genes referenced in prior scATAC-seq literature ^33^ and additionally collected marker genes known for their localization in specific cell types, as identified in other literature through *in situ* hybridization. We added the *ZmCLE7* gene from reference genome AGPv3.19 into the B73 v5 genome, by finding the sequence match at “chr4:8531149-8531929” for *ZmCLE7* in v5 using BLAST. To assess gene body chromatin accessibility, we counted Tn5 insertions within the gene body and 500 bp upstream and downstream extended regions using Granges and findOverlaps ^56^. Cell annotations were performed using a UMAP plot, depicting gene body accessibility per nucleus. For enhanced clarity in visualization, we applied smoothing to the normalized gene accessibility scores using a diffusion nearest neighbor graph ^57^. The other usage of gene body chromatin accessibility is to assess differences in gene body chromatin accessibility between genotypes. For this purpose, we calculated the log2 FC in chromatin accessibility by excluding genes with fewer than 50 Tn5 insertions in any cell type to mitigate extreme values resulting from sparse data.

#### Identification of accessible chromatin regions

Accessible chromatin regions were identified by aggregating cells with the same annotation into pseudo bulks. Peak calling was executed on pooled and individual replicates. MACS2 (v 2.2.7.1) was utilized with ‘--nomoel --keep-dup auto --extsize 150 --shift 75 --qvalue .05’ ^58^. The summits identified in each replicate were expanded by 250 bp on both sides. The peaks with less than 20 Tn5 coverage were filtered. Only the peaks present in all replicates and that overlapped a pooled set peak were retained ^51^.

#### Correlation analysis for the ACR by cell types

To compare chromatin accessibility within peaks across cell types, we used peaks called individually for each genotype and corresponding cell type. For the correlation analysis, we established a set of 500 bp union peaks. When using a fixed size of 500bp peaks, we assessed the p-value of each peak across cell types, utilizing a chromatin accessibility score that was normalized per million reads to identify the most representative peaks ^59^. The aggregated Tn5 insertions for each cell type were quantile normalized, and Pearson correlation coefficients were calculated using read density for the 2,000 most variable peaks.

#### Differential ACR analysis

For differential ACR analysis, we created the sets of union peaks per cell type. We merged peaks from both genotypes for each cell type, resulting in distinct sets of union peaks specific to each genotype. Consequently, the lengths of the peaks used for differential ACR analysis vary. Two pseudo replicates were generated for each biological replicate within cell types to enhance the robustness by randomly partitioning cells in one replicate into two groups. To quantify chromatin accessibility in the pseudo bulk replicates, we aggregated the chromatin accessibility within peaks by pseudo replicates to each cell type. We normalized aggregated chromatin accessibility as Trimmed Mean of M values (TMM) to adjust for different library sizes and performed statistical tests under generalized linear model using edgeR (v3.32.1)60.

#### Motif analysis in accessible chromatin regions

We used three approaches to find the motif enrichment in ACRs: 1) Motif deviation calculation across cells, 2) *de novo* motif searches and 3) known motif searches. To identify the cells where the motif is active, we computed motif deviation score using chromVar ^43^. We used Tn5 insertion counts at intergenic ACRs per cell as input for chromVar. For motif PWM, we used the non-redundant core plant PWM database from JASPAR2022 and PWMs derived from DAP-seq motif discovery. We applied smoothing to the bias-corrected motif deviations for each nucleus, integrating them into UMAP embedding for visualization, which is the same method used for visualizing gene body chromatin accessibility ^57^. Additionally, we rescaled the bias-corrected motif deviations to fit a color scale ranging from −1 to 1 across all motifs.

*De novo* motif searches in differential ACRs was performed using XSTREME version 5.5.3 within the MEME suite package (v5.5.0) ^61,62^. XSTREME leverages STEME that finds the enriched motif in the test set relative to control sets ^63^. While the test set is differential ACRs, control regions were used to determine the significance of motif occurrence compared to the background. To create the control set, we randomly selected the same number and length of ACRs from all ACRs, ensuring that they had a similar GC content ratio to the test set. To account for varied ACR lengths in the test set, we ensured the control set featured a comparable distribution of lengths. The motif search with STREME halts when it encounters a succession of motifs with p-values exceeding 0.05, applying “–thresh” option. We showcased motifs boasting an e-value below 1. Furthermore, utilizing the identified de novo motifs, fimo scrutinized specific ACR regions ^64^, pinpointing motif occurrences with p-values under 0.0001.

#### DAP-seq analysis and motif discovery

Raw reads were trimmed using trimmomatic ^65^ with the following settings: ILLUMINACLIP:TruSeq3-PE.fa:2:30:10:2:keepBothReads LEADING:3 TRAILING:3 SLIDINGWINDOW:4:15 MINLEN:50.

Trimmed reads were mapped to the B73v5 genome using bowtie2 ^66^ with default settings. Mapped reads were filtered to retain those with a MAPQ score greater than 30 using samtools ^67^ view -b -q 30. To identify peaks and motifs, we employed GEM (v 3.0) ^68^ with the following options: ‘--q 5 --k_min 5 -- k_max 14’. This option uses significant peaks with a q-value below 5 and specifies a range of k-mer lengths from 5 to 14 for motif discovery.

## Acknowledgements

This research was supported by the National Science Foundation (IOS-2026554) to RJS and AG as well as the UGA Office of Research to RJS.

## Author contributions

S.B., R.J.S., and A.G. designed the experiments and wrote the paper; S.B. and M.G. analyzed the data; X.Z., J.G., Z.C., M.G. and A.G. generated materials and/or conducted the experiments.

## Declaration of interests

R.J.S. is a co-founder of REquest Genomics, LLC, a company that provides epigenomic services. The remaining authors declare no competing interests.

## Supplemental information

Supplementary Tables.xlsx: TableS1-S5 in excel file.

Table S1. The quality information for the replicates.

Table S2. Meta data for the annotated cells of WT.

Table S3. Meta data for the annotated cells of *Bif3*.

Table S4. The number of peaks by cell types in WT and *Bif3*.

Table S5. Closet gene information for differential ACR and intergenic ACR in central zone.

Page 24-31: Figure S1-S8.

---

**Figure S1.**
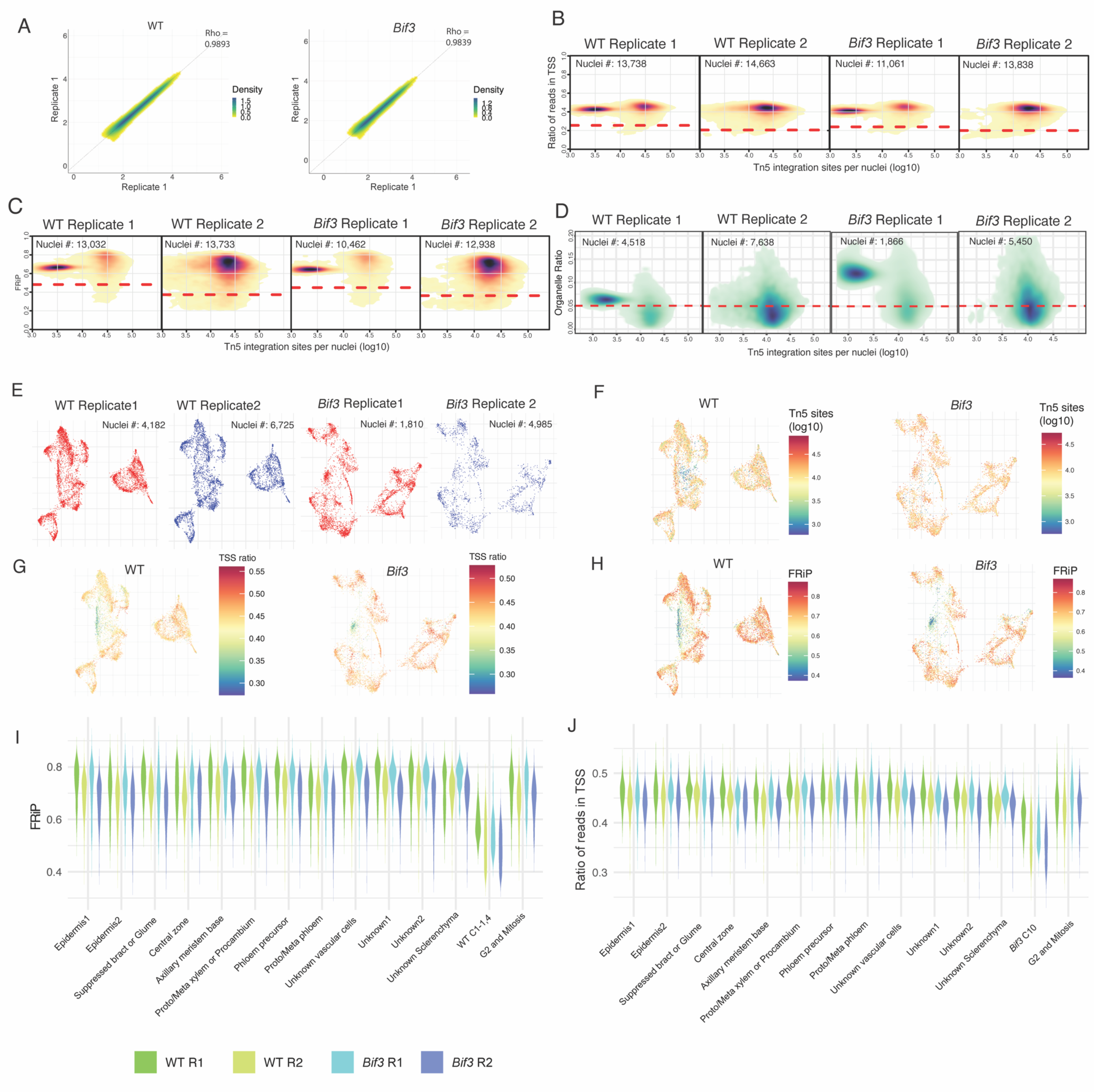
Evaluation and quality control of maize ear scATAC-seq libraries in WT and *Bif3*. **(A)** Correlation of replicates for the union peaks. The peaks were identified in bulk and the chromatin accessibility in peaks were quantile normalized using a log10 scale. **(B-D)** Density heatmap for ratio of reads in TSS, FRiP scores, organelle ratio by number of Tn5 per nuclei. Darker color represents dense nuclei number. The red line shows the cutoff for each library. **(E)** UMAP visualization for replicates. Replicates were well integrated by using UMAP dimensionality reduction of the Tn5 insertion sites after mitigating technical artifacts **(F-H)** The number of Tn5 insertions using a log10 scale, ratio of reads in TSS, and FRiP score by cell in UMAP. **(G, H)** FRiP and ratio of reads in TSS by annotated cell types. The four colors represent each of the libraries. **(I, J)** The FRiP score and ratio of reads in TSS of samples by cell types.

**Figure S2.**
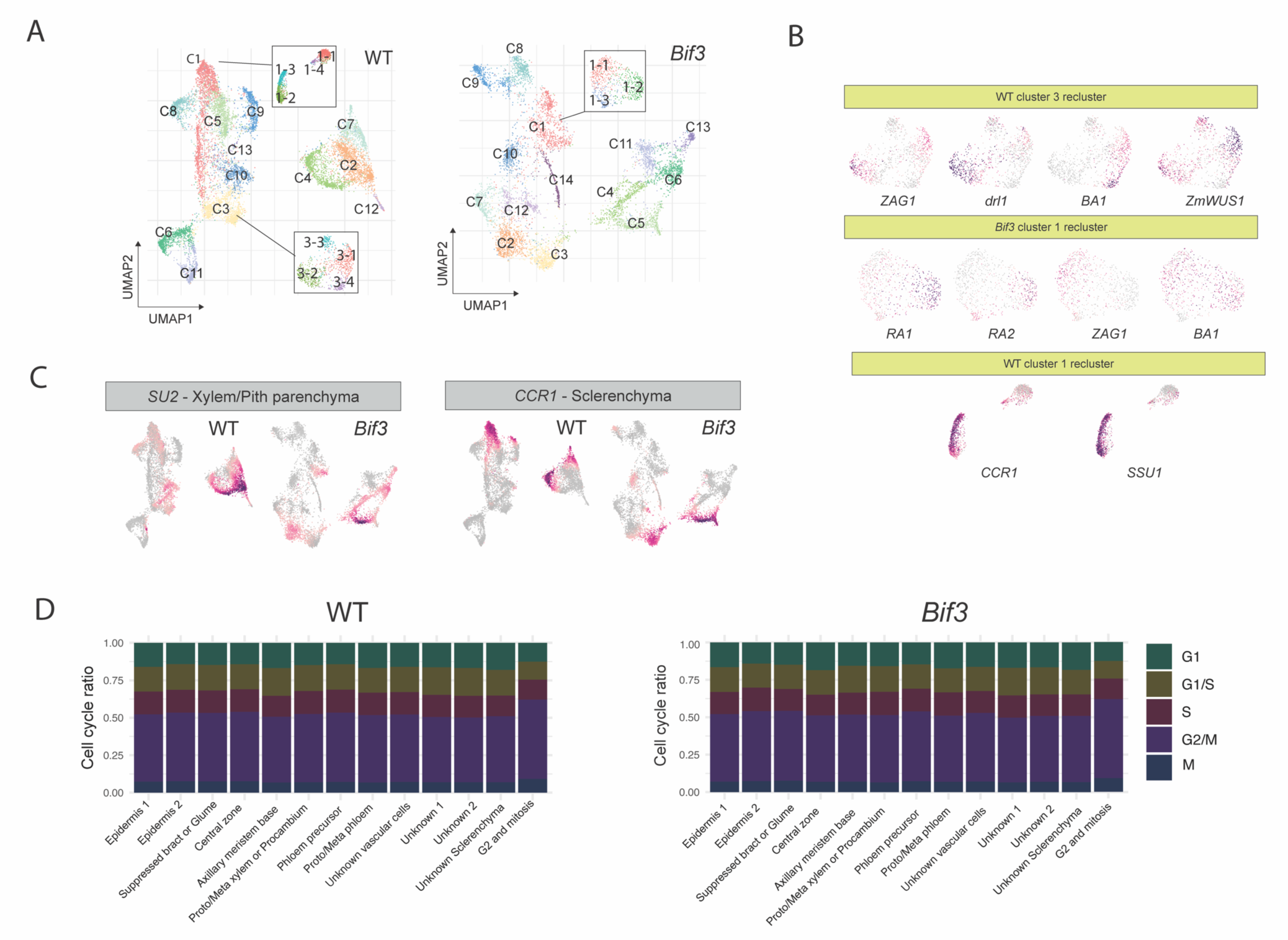
Annotation of clusters. **(A)** UMAP visualization representing WT and *Bif3*, categorized by cluster numbers. A highlighted inset focuses on the subclusters within WT C1, WT C3, and *Bif3* C8. **(B)** Gene body chromatin accessibility for specific marker genes across different subclusters: WT C3-1,2,3,4, *Bif3* C1-1,2,3 and WT C1-1,2,3,4. **(C)** The gene body chromatin accessibility for the ground cell marker genes. **(D)** Predicted the cell cycle phase by cell types in WT and *Bif3*. The colors represent the cell cycle phase for G1, G1/S, S, G2/M and M.

**Figure S3.**
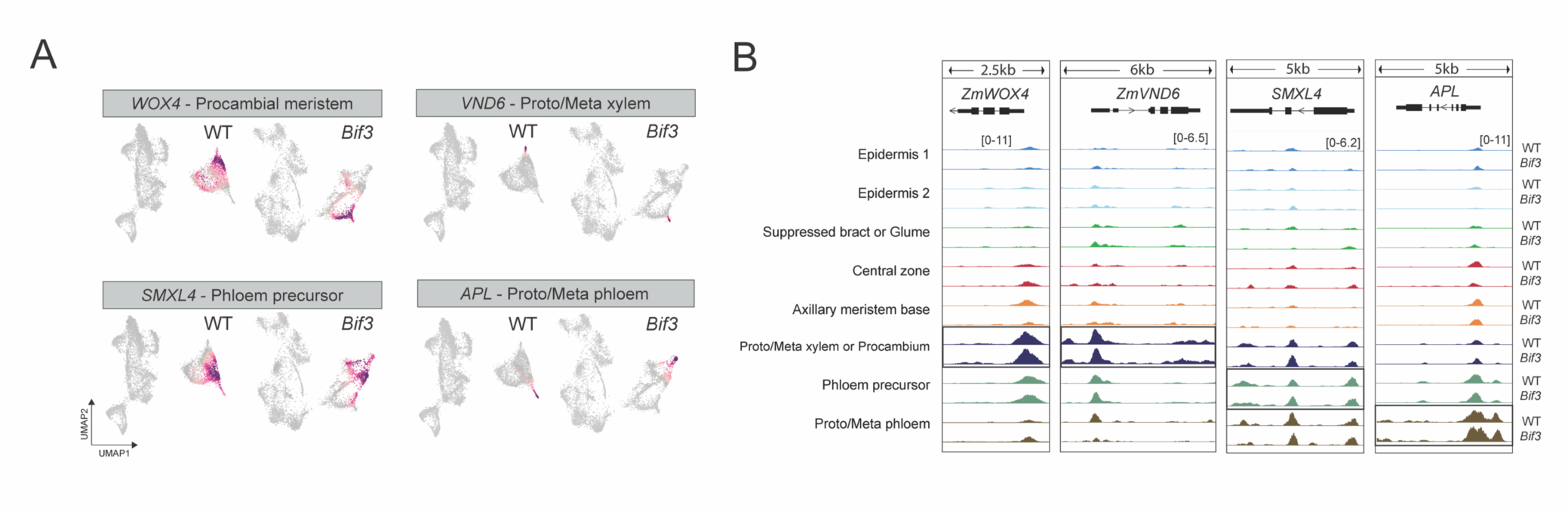
Annotation of vasculature cells. **(A)** Visualization of gene body chromatin accessibility for marker genes for vascular cells, with higher chromatin accessibility indicated in purple. **(B)** Genome browser tracks show chromatin accessibility by cell types around vascular cell marker genes in WT and *Bif3*.

**Figure S4.**
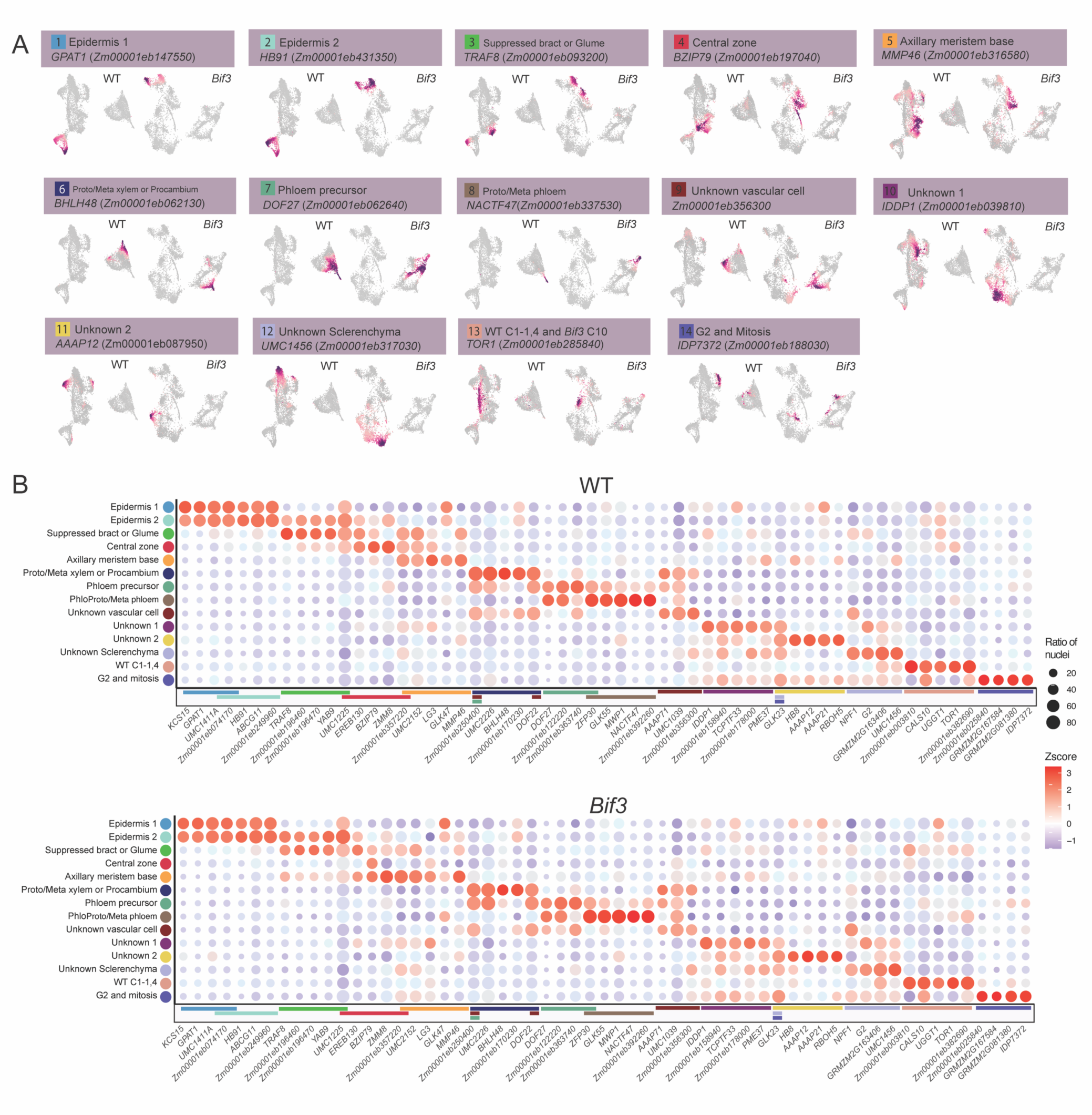
*De novo* marker genes identified across 14 clusters in WT. **(A)** UMAP highlighting areas of high chromatin accessibility, depicted in red, surrounding *de novo* marker genes in WT and *Bif3*. **(B)** Dot plot depicting five *de novo* marker genes for each cell type in WT and *Bif3*. The color above each *de novo* marker gene indicates the cell types where these genes exhibit higher gene body chromatin accessibility compared to other cell types. Five *de novo* marker genes were selected based on their statistical significance. Overlapping colors on the bars represent *de novo* marker genes that are shared between cell types.

**Figure S5.**
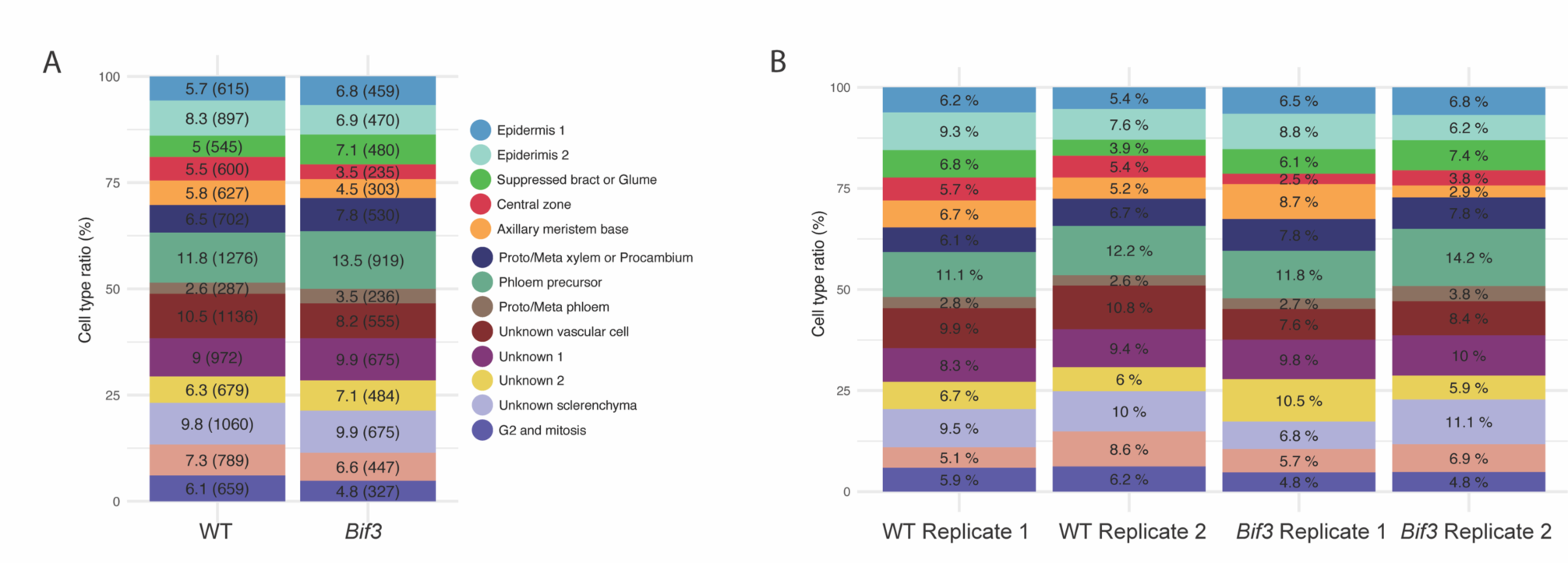
The distribution of cell types among the total cell population. **(A)** Proportion of each cell type in WT and *Bif3* samples. The numbers enclosed in parentheses indicate the number of cells. **(B)** Proportion of each cell type across libraries, with color coding consistent with the cell types in (A).

**Figure S6.**
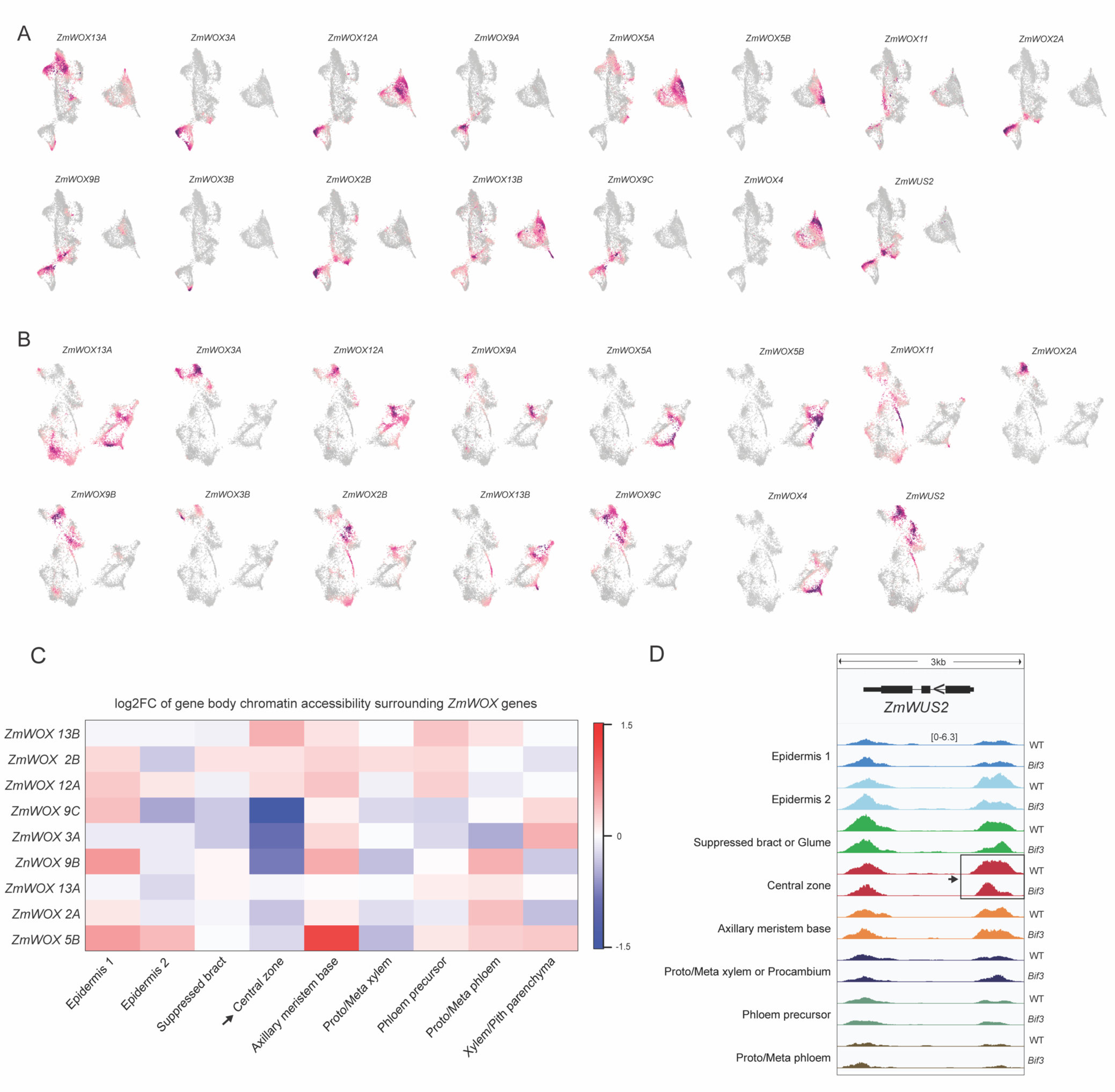
The gene body chromatin accessibility of *WOX* and *ZmWUS2* genes by cell types. **(A, B)** Visualization of gene body chromatin accessibility in UMAP of WT (A) and *Bif3* (B). **(C)** The log2FC of gene body chromatin accessibility for *WOX* genes between WT and *Bif3* mutant. **(D)** The genome browser image for chromatin accessibility around *ZmWUS2* gene by cell types and genotypes.

**Figure S7.**
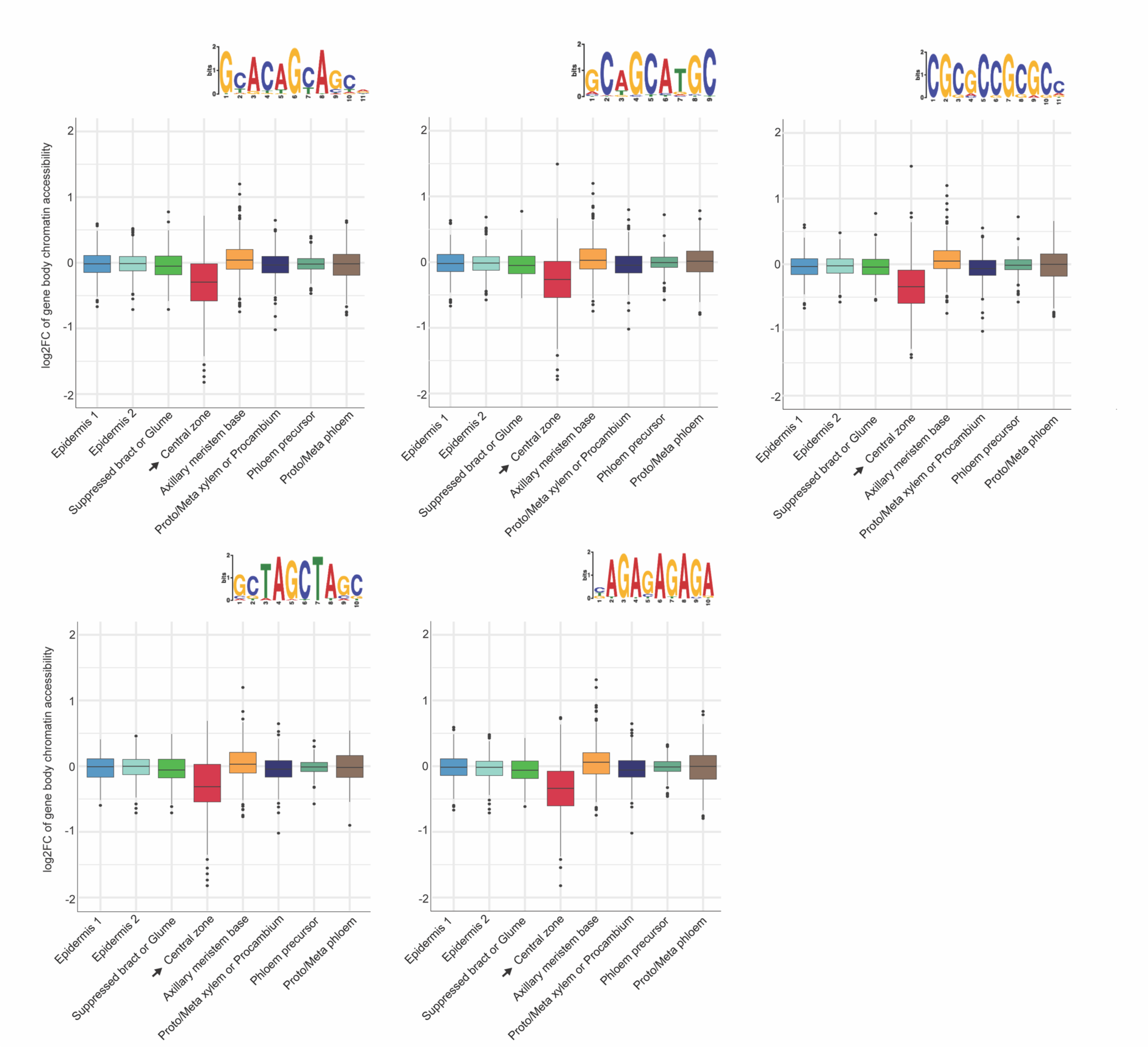
The boxplot illustrates the log2FC of gene body chromatin accessibility for genes located near decreased differential ACRs. Each plot represents a set of genes adjacent to decreased differential ACRs, categorized by the presence of specific motif sequences.

**Figure S8.**
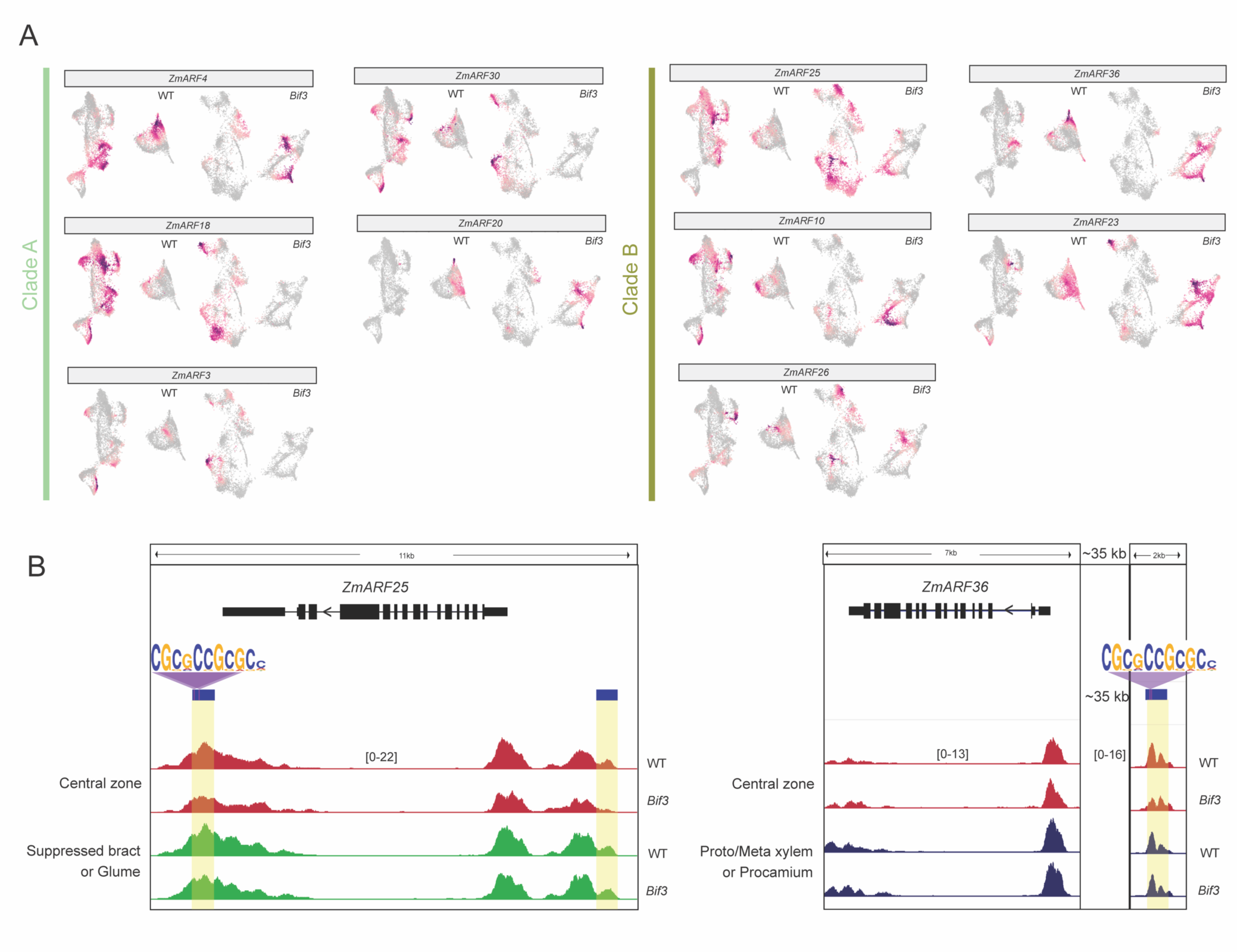
Chromatin accessibility of *ZmARF* genes in WT and *Bif3*. **(A)** UMAP visualization showing gene body chromatin accessibility for *ZmARF* genes, with genes categorized by clades. **(B)** Visualization of differentially accessible chromatin regions surrounding clade B *ZmARF* genes, highlighting the CGCGCCGCGCC motif presence in these differential ACRs. Blue bars and yellow highlights indicate the intergenic differential ACRs.

